# Inositol hexakisphosphate (IP6) and inositol pentakisphosphate (IP5) are required for viral particle release of retroviruses belonging to the primate lentivirus genus

**DOI:** 10.1101/2020.05.21.108167

**Authors:** Clifton L Ricana, Terri D Lyddon, Robert A Dick, Marc C Johnson

**Author notes:** **Corresponding Author:** 471C Bond Life Sciences Center, 1201 Rollins Street, Columbia, MO 65211.

## Abstract

Inositol hexakisphosphate (IP6) potently stimulates HIV-1 particle assembly *in vitro* and infectious particle production *in vivo*. However, knockout cells lacking the enzyme inositol-pentakisphosphate 2-kinase (IPPK-KO), which adds the final phosphate to inositol pentakisphosphate (IP5) to produce IP6, were still able to produce infectious HIV-1 particles at a greatly reduced rate. HIV-1 *in vitro* assembly can also be stimulated to a lesser extent with IP5, but it was not known if IP5 could also function in promoting assembly *in vivo*. IPPK-KO cells expressed no detectable IP6 but elevated IP5 levels and displayed a 20-100-fold reduction in infectious particle production, correlating with lost virus release. Transient transfection of an IPPK expression vector stimulated infectious particle production and release in IPPK-KOs but not in wildtype cells. Several attempts to make an IP6 and IP5 deficient stable cell line were not successful, but transient expression of multiple inositol polyphosphate phosphatase-1 (MINPP1) into IPPK-KOs resulted in the near ablation of IP6 and IP5. Under these conditions, HIV-1 infectious particle production and virus release were essentially abolished (1000-fold reduction). However, other retroviruses including a Gammaretrovirus, a Betaretrovirus, and two non-primate Lentiviruses displayed only a modest (3-fold) reduction in infectious particle production from IPPK-KOs and were not significantly altered by expression of IPPK or MINPP1. The only other retrovirus found that showed a clear IP6/IP5 dependence was the primate (macaque) Lentivirus Simian Immunodeficiency Virus (SIV-mac), which displayed similar sensitivity to IP6/IP5 levels as HIV-1. Finally, we found that loss of IP6/IP5 in viral target cells had no effect on permissiveness to HIV-1 infection. However, because it was not possible to generate viral particles devoid of IP6 and IP5, we were not able to determine if IP6 or IP5 derived from the virus producer cells is required at additional steps beyond assembly.

**Author Summary:** Inositol hexakisphosphate (IP6) is a co-factor required for efficient production of infectious HIV-1 particles. The HIV-1 structural protein Gag forms a hexagonal lattice structure. The negatively charged IP6 sits in the middle of the hexamer and stabilizes a ring of positively charged lysines. Previously described results show that depletion of IP6 reduces, but does not eliminate, infectious virus production. This depletion was achieved through knock-out of inositol-pentakisphosphate 2-kinase (IPPK-KO), the enzyme responsible for adding the sixth and final phosphate to the molecule. Whether IP6 is required, another inositol phosphate can substitute, or IP6 is simply acting as an enhancer for virus production was unknown. Here, we show that loss of IP6 and inositol pentakisphosphate (IP5) abolishes infectious HIV-1 production from cells. We do this through a cell-based system using transiently expressed proteins to restore or deplete IP6 and IP5 in the IPPK-KO cell line. We further show that the IP6 and IP5 requirement is a feature of primate lentiviruses, but not all retroviruses, and that IP6 and IP5 is required in the producer but not the target cell for HIV-1 infection.

## Introduction

The HIV-1 structural protein (Gag) is produced in the cytoplasm and traffics to the plasma membrane where it assembles into a viral particle that buds from the host membrane [1]. Gag is a polyprotein consisting of the Matrix (MA), Capsid (CA), Spacer 1 (SP1), Nucleocapsid (NC), Spacer 2 (SP2), and p6 domains [1,2]. During assembly, the Gag protein assembles into an ‘immature’ hexagonal lattice, driven primarily by interactions involving the CA and SP1 domains [1,3]. The C-terminal CA and SP1 domains contain an alpha-helix that forms a six-helix bundle with the other Gag proteins in the hexamer [4,5]. This bundle is important in formation and stabilization of the immature lattice [4,5]. During or shortly after budding from the cell, the viral protease cleaves the Gag polyprotein into its constitutive components, which separates CA from SP1 and eliminates the six-helix bundle [1,6]. The liberated CA protein then assembles into a structurally distinct ‘mature’ lattice which forms the viral core [1,7].

Early attempts to assemble full length HIV-1 Gag protein *in vitro* revealed that proper assembly required the presence of cell lysate [8]. This pointed to an assembly co-factor that catalyzed viral assembly in cells. Further research revealed that inositol phosphates were sufficient to stimulate proper assembly, but the mechanistic basis for this effect was poorly understood [3]. Recently, a Cryo-EM reconstruction of *in vivo*-produced immature HIV-1 particles revealed a small density above the CA-SP1 six-helix bundle that was coordinated by two rings of lysine residues, suggesting the presence of a negatively charged molecule inside the particle that helped stabilize the bundle [4]. This evidence for such a molecule, in conjunction with previous data that inositol phosphates stimulate assembly, prompted further evaluation of the role of inositol phosphates as HIV-1 assembly co-factors [3,8,9]. In particular, Inositol hexakisphosphate (I(1,2,3,4,5,6)P_6_ or IP6), which is a hexagonal six-carbon ring with a negatively charged phosphate at each position, seemed like a likely match for the density identified in particles [10].

In assembly experiments *in vitro*, the presence of IP6 was found to potently promote immature assembly and even to modulate whether particular Gag proteins assembled into immature or mature lattices [10]. Mutation of the lysine residues in Gag believed to coordinate the negatively charged molecule made the Gag proteins 100-fold less responsive to IP6 in *in vitro* assembly reactions [10]. When a crystal structure of the HIV-1 CA_CTD_SP1 protein in the presence of IP6 was solved, a density was observed associated with the six-helix bundle that precisely matched the density described in the Cryo-EM reconstruction [4,10]. These biochemical and structural data strongly support the conclusion that the density observed in HIV-1 particles is indeed IP6, but the data could not reveal whether IP6 is a requirement for HIV-1 assembly *in vivo.*

IP6 is found in mammalian cells at concentrations of 10-100uM [11], and is synthesized by a series of host enzymes through a complex and not fully resolved process (Fig 1A) [12–28]. The immediate precursor to IP6 is inositol pentakisphosphate (I(1,3,4,5,6)P_5_ or IP5), and the only enzyme known to catalyze the addition of the final 2-phosphate is inositol-pentakisphosphate 2-kinase (IPPK) [12–16]. IP5 was also shown to stimulate immature HIV-1 assembly *in vitro*, though not as robustly as IP6 [10,29–31]. In cells, several pathways have been described that lead to IP5 production, but all of those described in mammalian cells require the enzyme inositol-polyphosphate multikinase (IPMK) [12,16–20]. However, cells derived from a homozygous mouse embryo deficient in IPMK still produced residual levels of IP5 and IP6 through an unknown mechanism [15,17]. Recently, a genetic screen performed to identify genes involved in necroptosis revealed that inositol phosphates IP5 or IP6 are required for this process [21]. Importantly, the screen identified the genes *IPMK* and *inositol-tetrakisphosphate 1-kinase (ITPK1)* as being required for necroptosis, and cells lacking either of these genes were noticeably deficient in IP5 and IP6 [21,22]. Thus, ITPK1 and IPMK likely cooperate in the production of IP5 in cells. Knockouts of the IPPK and IPMK genes have both been reported to reduce the production of infectious HIV-1 particles *in vivo* [10,29–31].

**Fig 1.**
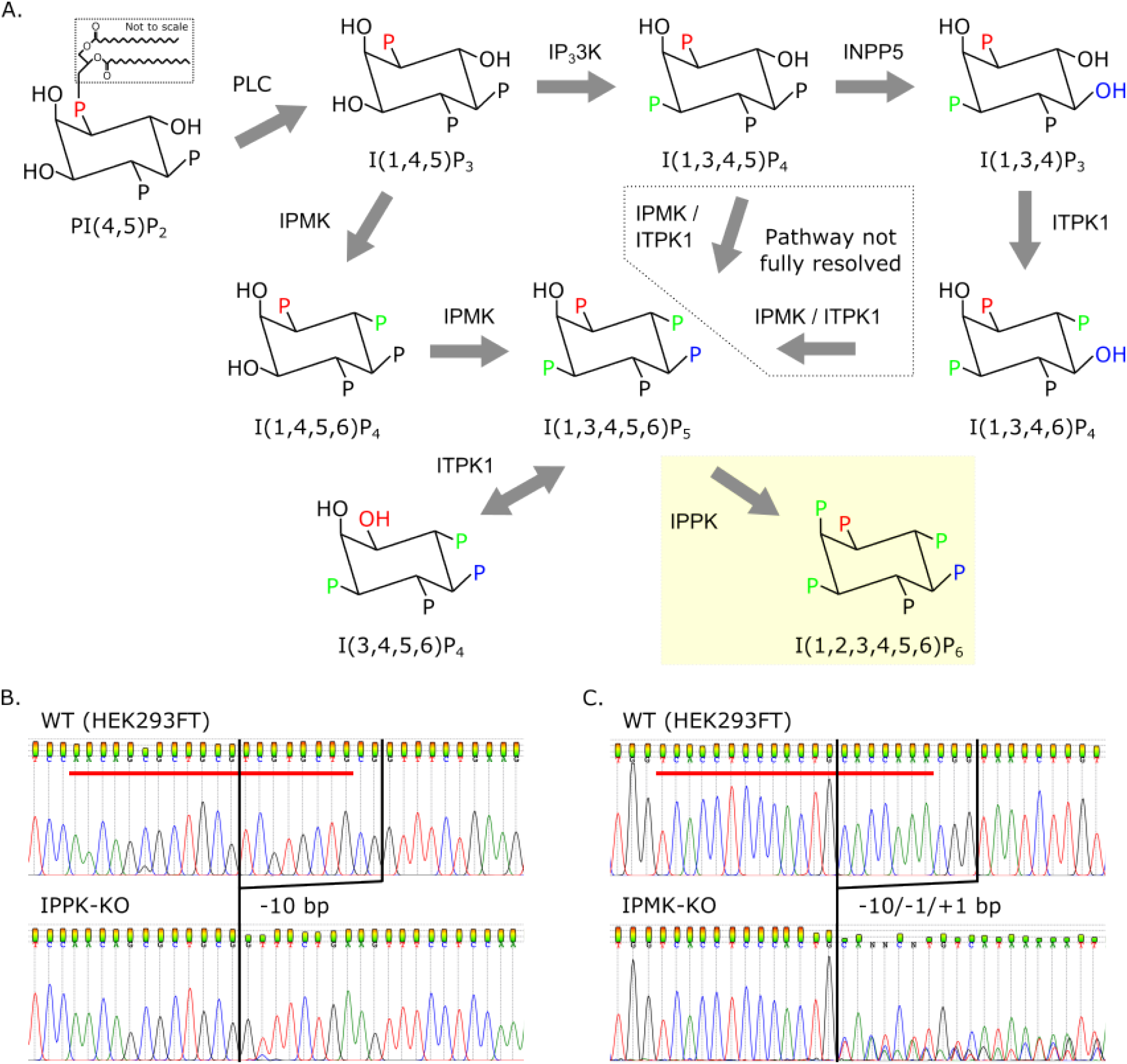
Knock-out of cellular genes leading to the production of IP6. (A) Inositol phosphate pathway in *H. sapiens*. Inositol-pentakisphosphate 2-kinase (IPPK) adds the sixth phosphate to position 2 of IP5 (yellow box). IP5 synthesis from I(1,3,4,5)P_4_ and I(1,3,4,6)P_4_ has not been fully resolved. Other abbreviations: phospholipase C (PLC), IP_3_3K (inositol-triphosphate 3-kinase), IPMK (inositol-polyphosphate multikinase), INPP5 (inositol-polyphosphate 5-phosphatase), and ITPK1 (inositol-tetrakisphosphate 1-kinase). (B-C) Chromatograms showing insertion-deletions of inositol-phosphate pathway KOs in HEK293FTs. Red bars delineate the 20-base pair guide RNA sequence used for CRISPR/Cas9 targeting. (B) KO of IPPK has a 10-base pair (bp) deletion. (C) KO of IPMK has three copies with 1- and 10-bp deletions and a 1-bp insertion.

Here, we sought to determine whether IP5 is required for the production of infectious HIV-1 particles in cells deficient in IP6. To accomplish this, we developed a system to transiently deplete cells of both IP5 and IP6 and test the assembly competence of HIV-1 and various other retroviruses under these conditions. We found that HIV-1 is dependent on the presence of IP5 or IP6 for viral production, but non-primate lentiviruses and viruses from retrovirus genera other than lentivirus are not. We further found that neither IP5 nor IP6 is required in viral target cells for successful HIV-1 infection.

## Results

### IPMK contributes to IP6 synthesis and infectious virus production

We previously showed that IP6 stimulates immature *in vitro* HIV-1 particle assembly [10]. We further showed that knock out of IPPK, the only enzyme known to catalyze addition of the final phosphate in the generation of IP6 (Fig 1A and B) [12–16], resulted in a drastic reduction in infectious HIV-1 viral particle production from cells [10]. However, infectious particles were still produced from HEK293FT IPPK knockout (IPPK-KO) cells, albeit at a greatly reduced rate [10]. There are three possible explanations for this partial phenotype. First, the cells could be continuing to produce low levels of IP6 through an unknown mechanism. Second, IP6 may enhance infectious HIV-1 particle production, but not strictly be required for it. Finally, IP5, the precursor to IP6 which can partially stimulate HIV-1 particle production *in vitro*, could substitute for IP6 in infectious particle production. To test the latter explanation, we attempted to generate a cell line that is deficient in both IP5 and IP6 production, in order to test if such a cell line would be completely deficient in infectious particle production. The enzyme inositol-polyphosphate multikinase (IPMK) has been reported to catalyze the penultimate step in IP6 production; thus, knockout of this enzyme would theoretically abolish IP5 and IP6 synthesis [12,16–22]. We used CRISPR/Cas9 with a guide RNA against *IPMK* [32,33] to generate a clonal IPMK knockout (IPMK-KO) cell line (Fig 1C). The validated IPMK-KO was then compared to HEK293FT and the previously described IPPK-KO cell line (previously described in [10] and sequence validation shown in Fig 1B).

First, we wanted to validate the loss of IP6 and IP5 in our IPPK-KO and IPMK-KO cells. Using TiO_2_ extraction and 33% PAGE separation [34,35], we found that the IPPK-KO cells had no detectable IP6, but a slightly elevated level of IP5 compared to HEK293FT cells (Fig 2A-C). In contrast, the IPMK-KO cells had residual levels of both IP6 and IP5 (Fig 2A-C). This was consistent with a previous report that showed that cells from an IPMK knockout embryo were also shown to produce very low levels of IP5 and IP6 through an unknown mechanism [21,22].

**Fig 2.**
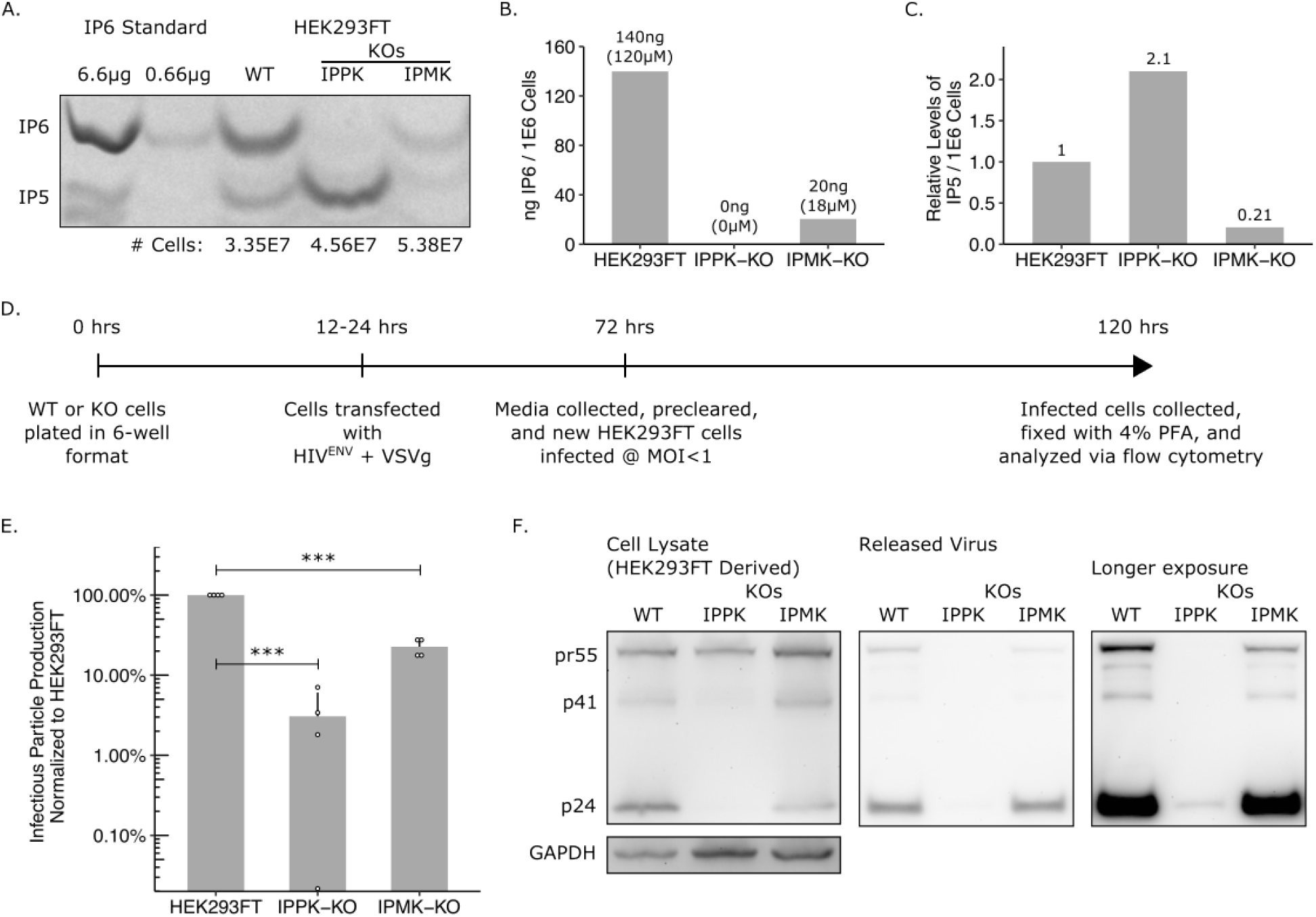
IP pathway KOs have reduced IP6 and IP5 levels and have a loss of infectious particle release. (A) 33% PAGE gel separating inositol phosphates. Two dilutions of purified 1 M IP6 were used as a standard and had IP5 breakdown products. The number of cells in each sample is indicated. (B) IP6 quantification of panel A in ng per million cells and μM. (C) Relative IP5 quantification normalized to the HEK293FT control. (D) Experimental timeline. (E) Percent infectious particle release normalized to HEK293FT cells. Student’s t-test was used for pair-wise comparison (n = 4, *** p < 0.001, error bars = mean + SD). (F) Representative western blot of experiments from panel D. Full-length HIV-Gag (pr55) and GAPDH loading control are presented on the left panels. Virus released into media is presented in the middle panel. A longer exposure of the virus release blot was also taken and presented on the right panel.

Next, we measured infectious HIV-1 virus particle production from HEK293FT cells and its two derivative knockout lines. An HIV-1^ΔEnv^ provirus containing a GFP reporter (HIV-CMV-GFP) was co-transfected with a VSV-G expression construct into the three cell lines in parallel, and the media was titered on fresh target cells (Fig 2D). As we reported previously, infectious particle production from the IPPK-KO cell line was reduced ~20-100 fold compared to HEK293FT cells (Fig 2E) [10]. The IPMK-KO cells displayed a more modest ~5-fold reduction in infectious particle production that corresponded with the residual IP6 levels found in the cells (Fig 2E). We then tested whether the block in infectious virus particle production was due to a block in virus release or to reduced infectivity of released virus. To accomplish this, we measured the Gag/CA protein level in producer cells and the supernatant. Western blots with an antibody against p24 CA revealed that HEK293FT, IPPK-KO, and IPMK-KO cells all produced full length Gag at relatively equal levels (Fig 2F, left). However, p24 CA released into the supernatant was barely detectable from the IPPK-KO cells, while media from IPMK-KO cells contained normal p24 levels (Fig 2F, middle and right). Together, the western blot data show that loss in infectious particle production primarily correlates with a loss in viral release.

### Exogenous addition of MINPP1 depletes IP6 and IP5 and is toxic to cells

Knock-out of IPPK ablates IP6 in cells while increasing IP5 levels, while knock-out of IPMK leaves residual levels of IP6 and IP5. The finding that low-level infectious particle production still occurs in IPPK-KO cells is consistent with the hypothesis that IP5 can weakly substitute for IP6 in supporting HIV-1 assembly. However, because neither IPPK-KO nor IPMK-KO cells were completely devoid of IP5 and IP6, we could not rule out other explanations. Therefore, we next attempted to make pairwise knockouts in IPPK, IPMK, and ITPK1 (which also contributes to IP5 synthesis [12,16,21–27]) to further reduce inositol phosphate levels. Numerous attempts failed to yield such double knockouts, likely because loss of the combination of enzymes was lethal. Therefore, as an alternative approach, we attempted to modulate levels of inositol phosphates by overexpressing multiple inositol polyphosphate phosphatase-1 (MINPP1), an enzyme that removes the 3-phosphate from IP6 and IP5 (Fig 3A) [12,36–38]. While removal of 3-phosphate from IP6 results in the production of an alternative species of IP5 (I(1,2,4,5,6)P_5_), this IP5 has an equatorial hydroxyl group which is likely not favorable for interaction with the lysines in the IP6 binding pocket (Fig 3A).

**Fig 3.**
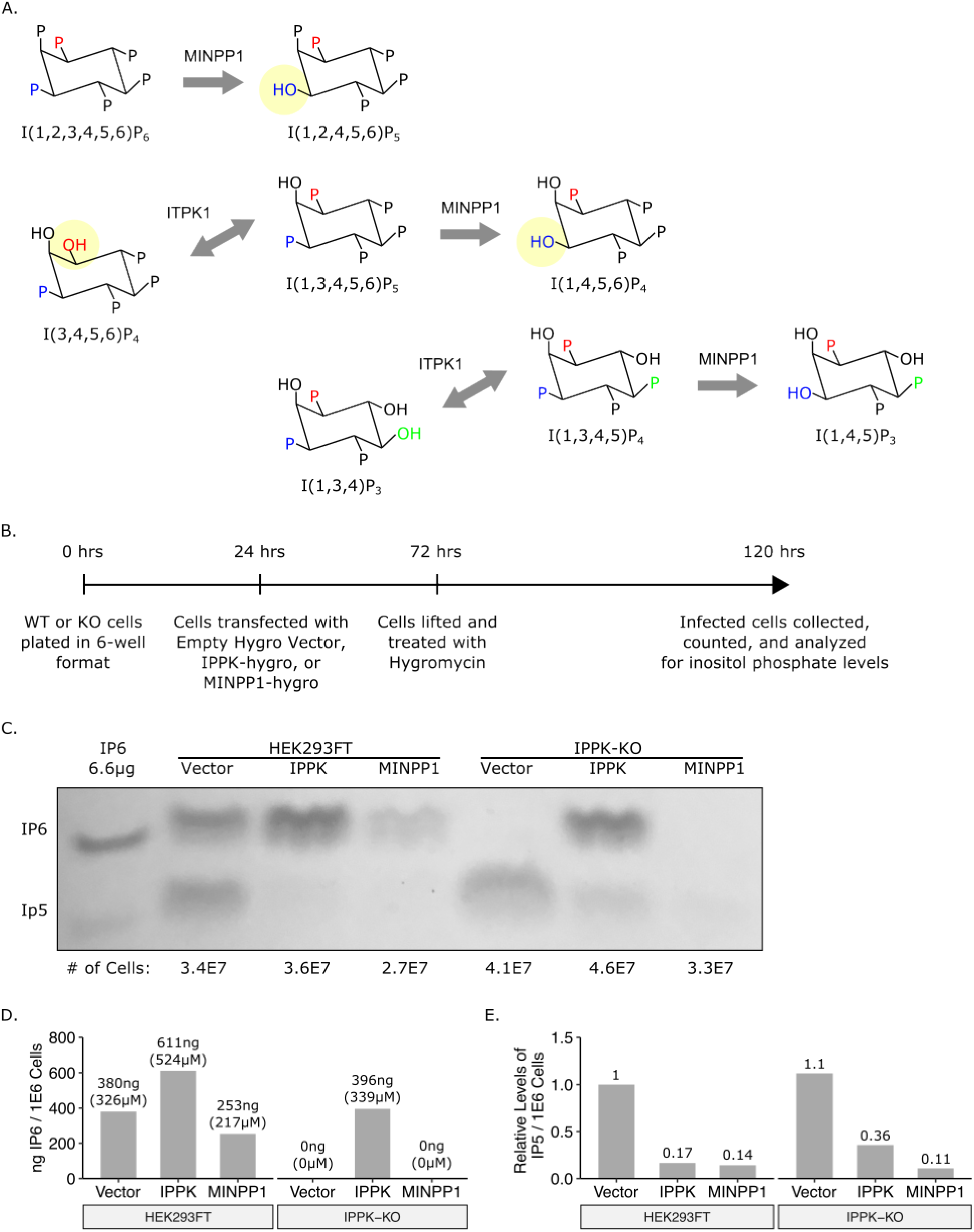
Addition of multiple inositol polyphosphate phosphatase 1 (MINPP1) removes endogenous IP6 and relevant IP5 species from cells. (A) Inositol phosphate pathway showing MINPP1 removal of the 3-position phosphate from IP6, IP5, and IP4. Removal of 3-phosphate from IP6 and I(1,3,4,5,6)P_5_ results in an equatorial hydroxyl group. (B) Experimental timeline. (C) 33% PAGE gel separating inositol phosphates. Dilution of purified 1 M IP6 was used as a standard and had an IP5 breakdown product. The number of cells in each sample is indicated. (C) IP6 quantification of panel C in ng per million cells and μM. (D) Relative IP5 quantification of panel C normalized to the HEK293FT control.

We first tested whether inositol phosphates could be modulated in a transient assay. To do this, we generated expression constructs containing cDNAs for IPPK or MINPP1 on a plasmid that also expressed a selectable hygromycin gene. These vectors were individually transfected into HEK293FT cells or IPPK-KO cells, the cells were briefly treated with hygromycin to eliminate untransfected cells, and inositol phosphate levels were directly measured from the surviving cells (Fig 3B). In HEK293FT cells, exogenous expression of IPPK increased IP6 levels while decreasing IP5 levels (Fig 3C-E, column 2). In contrast, exogenous expression of MINPP1 reduced, but did not ablate, IP6 and IP5 levels (Fig 3C-E, column 3). IPPK-KO cells had no detectable IP6, but expression of IPPK restored IP6 levels (Fig 3C-E, columns 4-5). Expression of MINPP1 in IPPK-KO cells resulted in a near complete loss of IP5 in addition to the loss of IP6 (Fig 3C-E, column 6). This result demonstrates that exogenous expression of MINPP1 is a viable method of modulating intracellular IP5 levels.

To reduce the inherent variability in transient transfection experiments, we attempted to stably express MINPP1 in IPPK-KO cells. These attempts to stably express MINPP1 resulted in poor recovery under hygromycin in IPPK-KO cells but was tolerated in HEK293FT cells. Previous reports indicated that MINPP1 expression can induce apoptosis [36–38]. To verify that the low recovery of MINPP1 expressing cells was in fact due to toxicity, we created a retroviral reporter vector in which MINPP1 cDNA was followed by an IRES-EGFP (Fig 4A). Cells transduced with this reporter should express both MINPP1 and EGFP, and cell survival can be determined by measuring the number of GFP positive cells over time (Fig 4B). HEK293FT cells expressing MINPP1-IRES-GFP or a control (IRES-GFP alone, Fig 4A) were maintained in the population over the course of three weeks indicating tolerance of MINPP1 (Fig 4C-D). However, the majority of IPPK-KO cells expressing of MINPP1-GFP were lost over the course of the experiment, consistent with toxicity (Fig 4C-D). This supports previous reports of a balance of IP6 and IP5 promoting cell viability [37,38]. Importantly, cell death from MINPP1 expression in IPPK-KO cells was not immediate, which allowed a window for testing the effects of MINPP1 expression on virus production.

**Fig 4.**
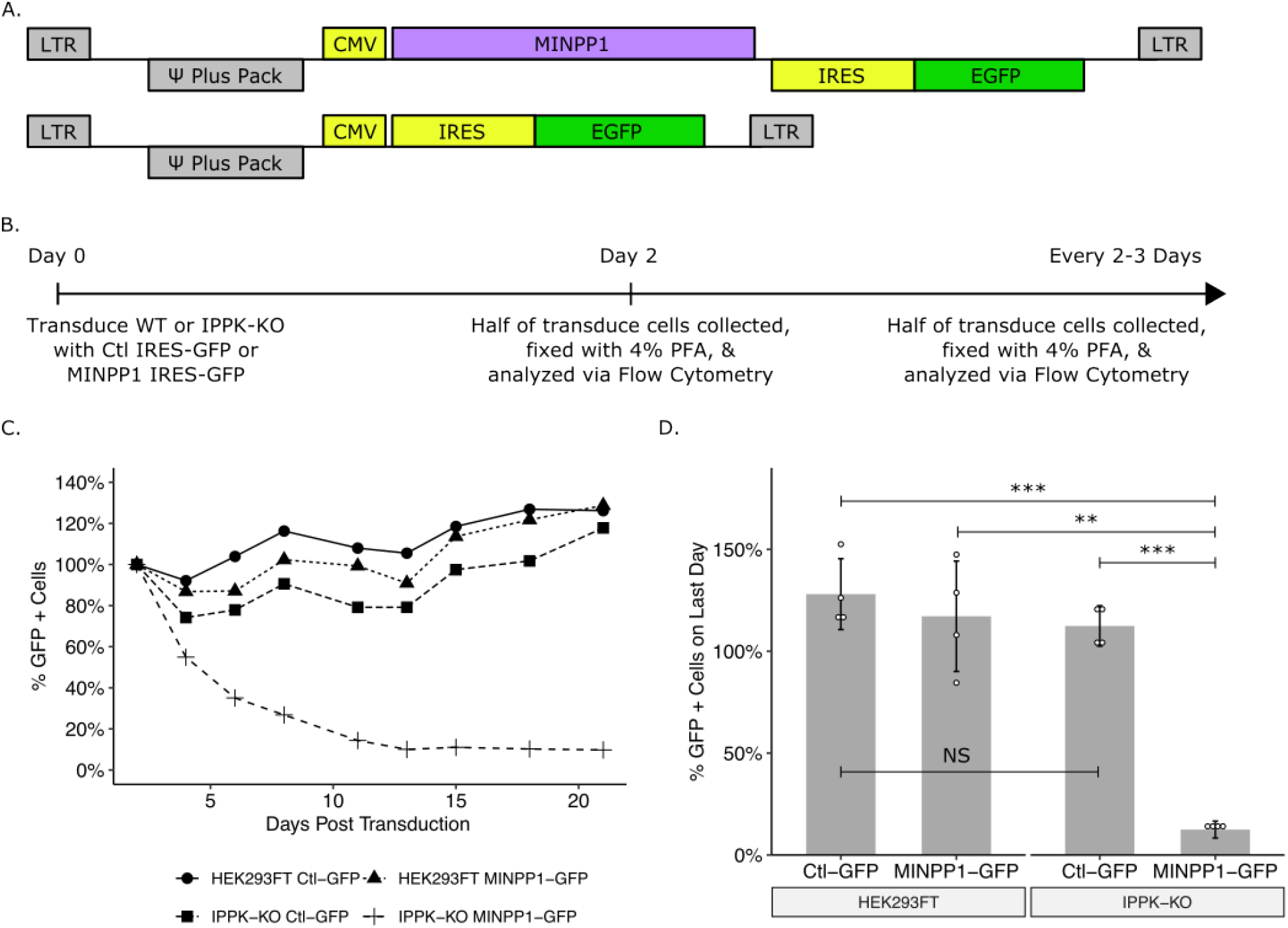
Exogenous expression of MINPP1 is toxic in IPPK-KO cells. (A) Plasmid map of expression vectors. (B) Experimental timeline. (C) Line plot of a representative experiment. The percentage of cells expressing EGFP over time are normalized to the starting population of EGFP positive cells for each cell line. (D) Bar chart of percent EGFP positive cells on the last day of collection from panel C (day 21). Student’s t-test was used for pair-wise comparison (n = 4, ** p < 0.01, *** p < 0.001, error bars = mean ± SD).

### Exogenous addition of MINPP1 results in near abolishment of HIV-1 infectious virus production

Because it was not possible to make a stable IPPK-KO cell line expressing MINPP1, we chose to test the effects of MINPP1 expression on virus production using a transient expression assay. Briefly, HIV-CMV-GFP and VSV-G DNAs were co-transfected into HEK293FT or IPPK-KO cells with either an expression vector containing IPPK cDNA, MINPP1 cDNA, or no insert, and the cells were allowed to produce virus for two days (Fig 5A). Infectious virus particles were then titered on HEK293FT cells (Fig 5A). As before, infectivity was reduced 20-100-fold from IPPK-KO cells (Fig 5B). Addition of IPPK to HEK293FT cells had no appreciable effect on this number, but addition to IPPK-KO cells enhanced infectious particle production by approximately 10-fold (Fig 5B). Likewise, addition of MINPP1 to HEK293FT cells also had no appreciable effect on infectious particle production, but addition to IPPK-KO cells further reduced infectivity by approximately 10-fold, which approached background levels (Fig 5B). Western blotting was again used to determine at which step in infectious virus particle production was blocked (Fig 5C). The expression of Gag in cells was similar among all conditions, indicating that modulation of inositol phosphate levels was not grossly affecting translation levels (Fig 5C, top and middle). Exogenous expression of IPPK and MINPP1 did not appear to affect viral release or protein maturation from HEK293FT cells (Fig 5C, bottom left). However, viral release from IPPK-KO cells varied considerably across the conditions, and the amount of virus released closely tracked with infectious particle production (Fig 5C, bottom right). HIV-1 CA release in IPPK-KO cells was greatly stimulated by expression of IPPK cDNA, but CA release was essentially abolished by expression of MINPP1. These data suggest that IP5 or IP6 is required for the release of viral particles. However, because there were essentially no virus particles released in the absence of IP5 and IP6, it was not possible to determine if any such particles might have been infectious. Thus, it remains possible that IP5 and/or IP6 also are required at other stages of the viral life cycle.

**Fig 5.**
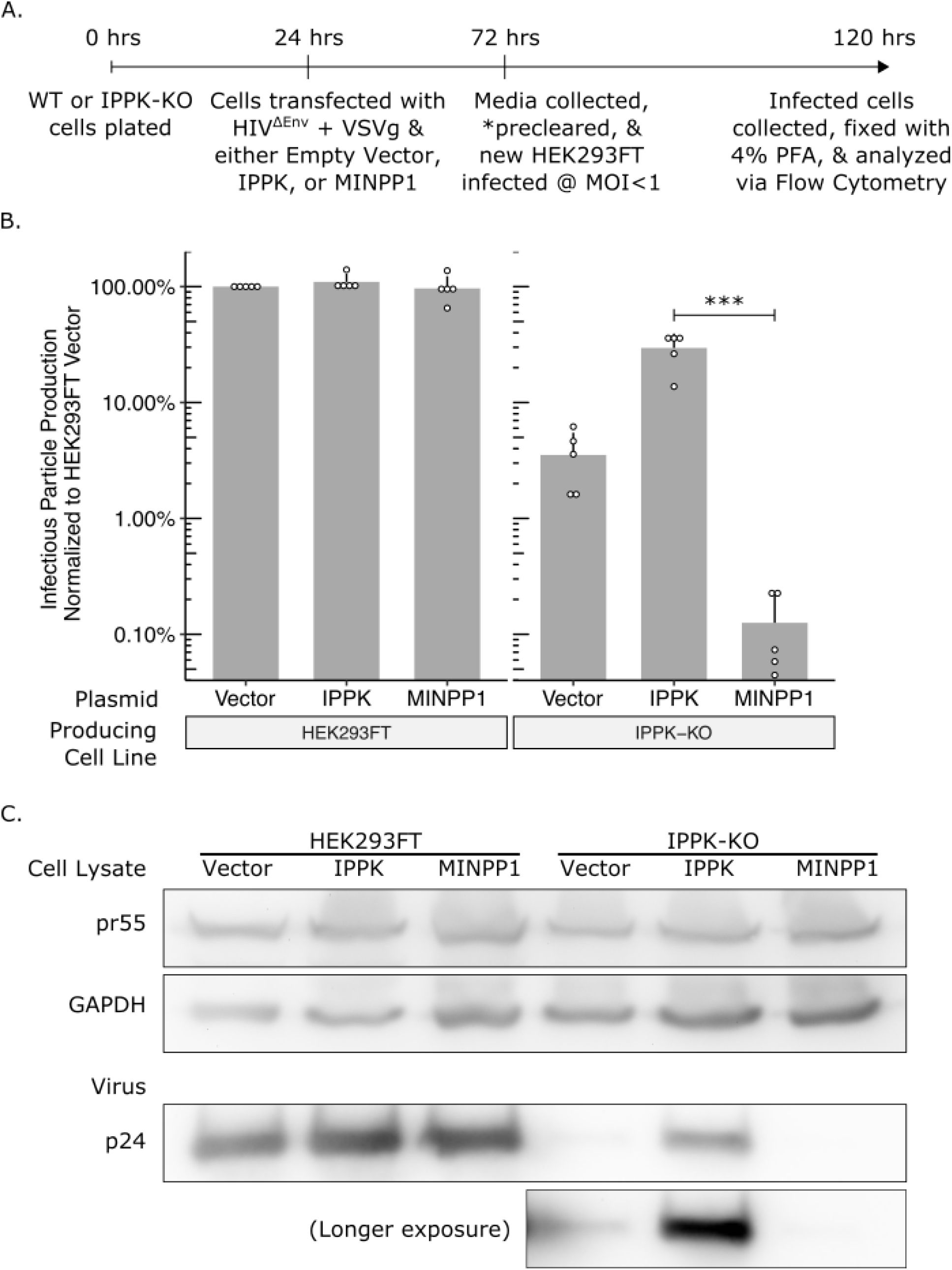
IPPK-KO cells expressing MINPP1 have substantial loss in infectious particle production due to a block in viral release. (A) Experimental timeline. (B) Bar chart of percent infectious particle release normalized to virus from HEK293FT cells expressing the empty vector. Student’s t-test was used for pair-wise comparison (n = 5, *** p < 0.001, error bars = mean ± SD). (C) Representative western blot of experiments from panel B. The rows are full-length HIV-Gag (top), GAPDH loading control (middle), and virus released into media (bottom). A longer exposure was also taken for the blot of released virus.

### Depletion of IP6 and IP5 in target cells does not affect susceptibility to HIV-1 infection

IP6 has been shown to stabilize the lattice of the immature and mature hexamer [10,29–31]. This stabilization has been proposed to be important for DNA synthesis following the release of the viral core into the cytoplasm of target cells [29]. When the viral core is depleted of IP6 *in vitro*, it breaks down more readily. Thus it has been inferred that after fusion with the target cell, IP6/IP5-lacking virus cannot effectively reverse transcribe the viral RNA to DNA [29,39]. To test if IP6 in the target cell is required for susceptibility to infection, we infected HEK293FT or IPPK-KO cells with virus produced from either HEK293FT or IPPK-KO cells and compared infection levels. If IP6 in the target cell is required for viral infection, one would expect to see a lower viral titer in IPPK-KO cells, regardless of the source of virus. By contrast, if IP6 is required for infection but can be derived from the producer cells, one would expect the virus to have a lower relative titer on IPPK-KO cells only when the virus is produced from IPPK-KO cells. In fact, we observed no significant difference in titer on the two types of cells, regardless of the source of the virus (Fig 6A). While this experiment demonstrates that IP6 is not required in the target cell for infection, it does not address whether IP5 is perhaps able to substitute for IP6 at this stage of the infection.

**Fig 6.**
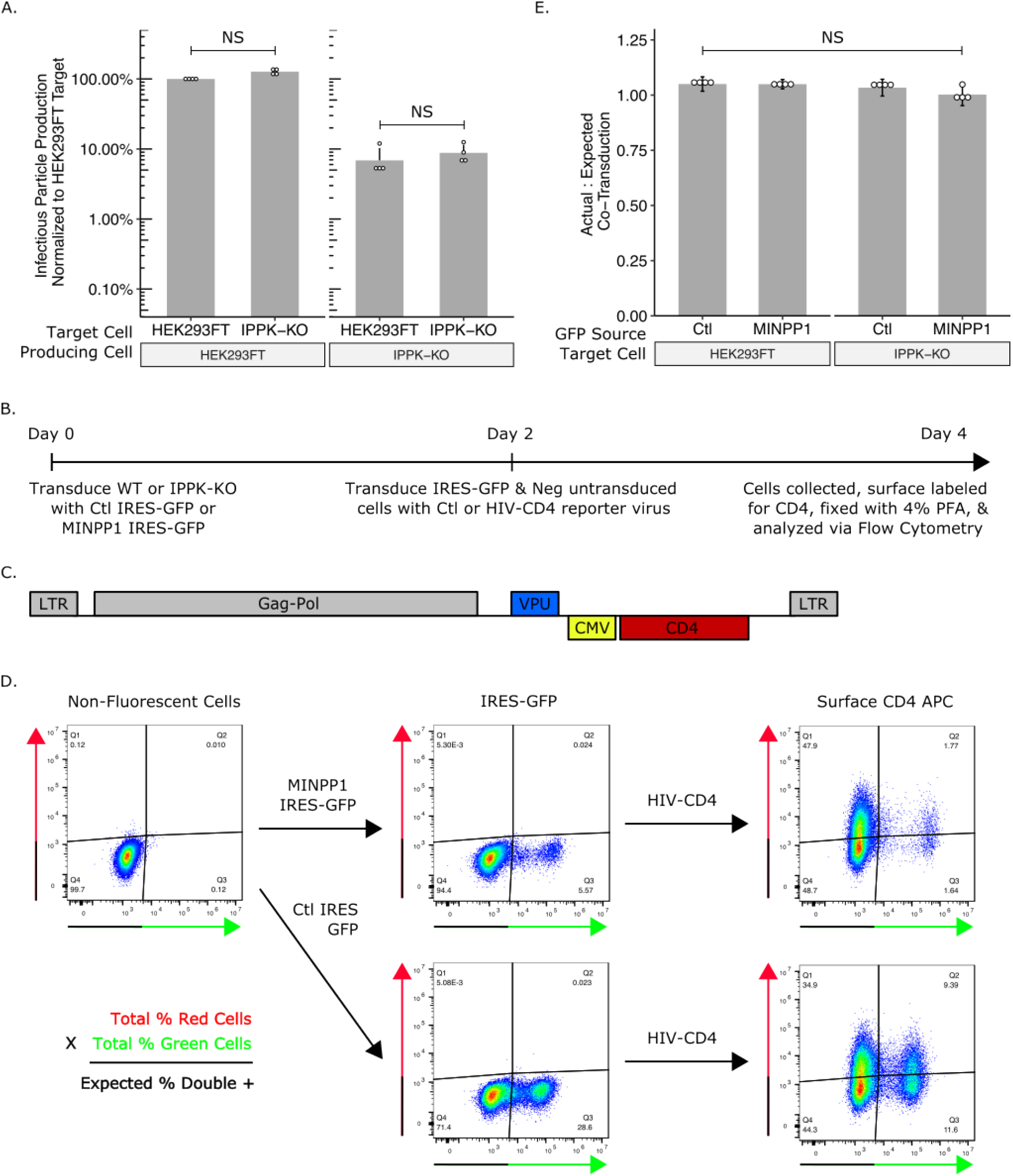
IP6 and IP5 levels in target cells do not affect susceptibility to HIV-1 infection. (A) Bar chart of percent infectious particle release normalized to HEK293FT cells. Student’s t-test was used for pair-wise comparison (n = 4, * p < 0.05). (B) Experimental timeline of the assay. (C) Plasmid map of HIV-1^ΔEnv^-CD4. (D) Example flow plots show output from the assay. (E) Bar chart of the ratio of the actual percent double positive cells to the expected double positive cells. The expected percent of double positive cells was calculated from the total percent of red cells and green cells. Student’s t-test was used for pair-wise comparison (n = 4, * p < 0.05, error bars = mean ± SD).

To test the role of IP5 depletion in susceptibility of target cells, we next transduced the MINPP1-IRES-GFP vector or an empty IRES-GFP control (Fig 4A) into HEK293FT or IPPK-KO cells, and then tested their susceptibility to infection (Fig 6B). Two days after transduction with MINPP1 IRES-GFP or IRES-GFP, cells were transduced with VSV-G pseudotyped HIV-1^ΔEnv^ virus containing a CD4 reporter (Fig 6C). Two days after virus transduction, surface CD4 was stained with an APC-conjugated antibody to score for successful virus transduction (Fig 6D). If cells expressing MINPP1 are less susceptible to infection, then the fraction of GFP-positive cells that are also APC positive (virus transduced) should be less than the fraction GFP-negative cells that are APC positive. If they are more susceptible, then the fraction of GFP-positive cells that are also APC positive should be more than the fraction GFP-negative cells that are APC positive. The expected fraction of GFP/APC double positive cells can then be calculated based on the total number of GFP and APC positive cell and compared to the actual number of GFP/APC double positive cells observed (Fig 6D, equation). Expression of MINPP1 was found not to alter the susceptibility of HEK293FT or IPPK-KO cells (Fig 6E). Together, these data suggest that neither IP6 nor IP5 from target cells is required for viral infection. As before, since we were not able to obtain virus that was devoid of IP5 and IP6, it was not possible to determine if IP5 and/or IP6 from the producer cell is required for core stability during infection.

### Beta- and Gamma-retroviruses do not require IP6 or IP5 for infectious virus production

Different retroviral species vary in Gag lattice structure and viral protein trafficking. The IP6 and IP5 requirement for assembly of other retroviral species can inform the different assembly strategies utilized by retroviruses. With HIV-1 as the model virus, we first tested outgroup retroviral species with our exogenous gene co-transfection system. With expression of IPPK and MINPP1 in HEK293FT cells, the Gammaretrovirus Murine Leukemia Virus (MLV) did not vary in viral output (Fig 7A). Infectious MLV particle production from IPPK-KO was reduced a few-fold compared to HEK293FT cells; however, this reduction could not be modulated further by addition of IPPK or MINPP1. Expression of IPPK actually caused a small but statistically insignificant reduction in infectious particle production compared to empty vector (Fig 7A). With expression of MINPP1 in IPPK-KO cells, there was no difference in virus output compared to empty vector (Fig 7A). These data suggest that the 3-fold reduction in virus particle release with MLV does not reflect a direct IP6 or IP5 requirement.

**Fig 7.**
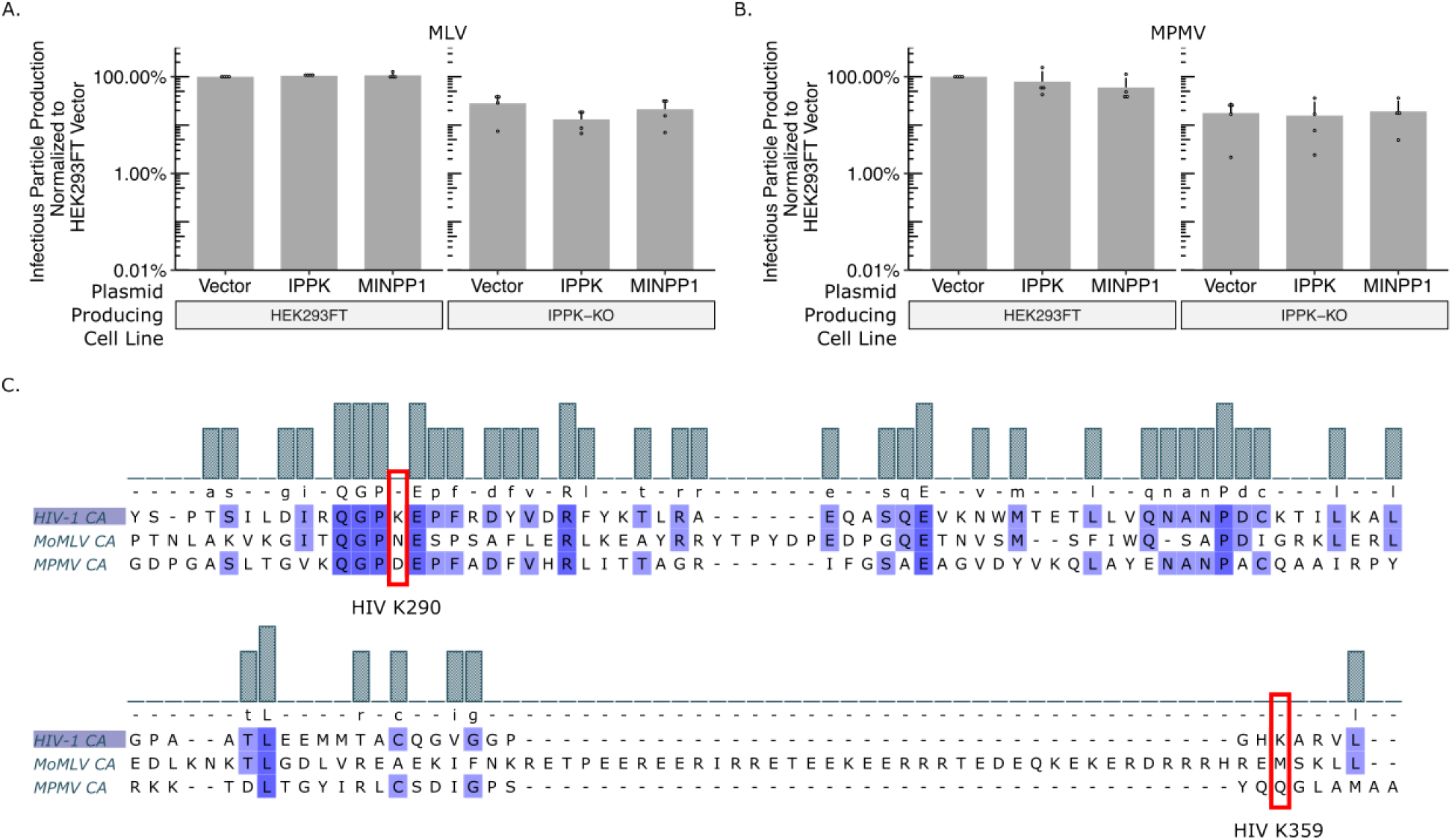
Gammaretroviruses and Betaretroviruses do not require IP6 or IP5 as assembly co-factors. Bar charts of percent infectious particle release of other retroviral genera normalized to virus from HEK293FT cells expressing the empty vector. (A) The Gammaretrovirus Murine leukemia virus (MLV, n = 4). (B) The Betaretrovirus Mason-Pfizer monkey virus (MPMV, n = 5). (C) Multiple sequence alignment of CA proteins of HIV-1, MLV, and MPMV. Note the lack of K290 and K359 homology in MLV and MPMV.

We next tested the Betaretrovirus Mason-Pfizer Monkey Virus (MPMV). As with HIV-1 and MLV, expression of IPPK and MINPP1 in HEK293FT cells did not affect infectious particle release (Fig 7B). Infectious particle production was again slightly decreased from IPPK-KO cells, but neither IPPK nor MINPP1 expression altered this output (Fig 7B). As with MLV, these data suggest that MPMV does not have a strict IP6 or IP5 requirement for infectious particle production. Additionally, amino acid sequence alignment between HIV-1, MLV, and MPMV CA proteins shows no homology to the K290 and K359 residues that interact with IP6 and IP5 in HIV-1 (Fig 7C) [10].

### The IP6 and IP5 requirement is conserved across primate lentiviruses

Both outgroups tested are from different retroviral genera than HIV-1. To determine whether IP6 and IP5 are required for other members of the Lentivirus genus, we next tested the primate (macaque) Simian Immunodeficiency Virus (SIV-mac), the feline Feline Immunodeficiency Virus (FIV), and the equine Equine Infectious Anemia Virus (EIAV). In our exogenous gene expression system, SIV had similar outputs to HIV-1 (Fig 8A). Infectious particle production was reduced over 20-fold from IPPK-KO cells relative to HEK293FT cells. Importantly, infectious particle production was partially restored with addition of IPPK and precipitously reduced by the addition of MINPP1. As with MLV and MPMV, transfection of IPPK-KO with EIAV and FIV produced about 3-fold fewer infectious virus particles than HEK293FT cells (Fig 8B-C). However, neither virus was significantly affected by introduction of IPPK or MINPP1 (Fig 8B-C). Comparison of the protein sequence alignments shows homology between all four lentiviruses at K290 and K359; however, prolines upstream and downstream of K290 are not conserved for FIV and EIAV (Fig 8D). Together, these data point toward an IP6 and IP5 requirement for primate lentiviruses but not lentiviruses of other species.

**Fig 8.**
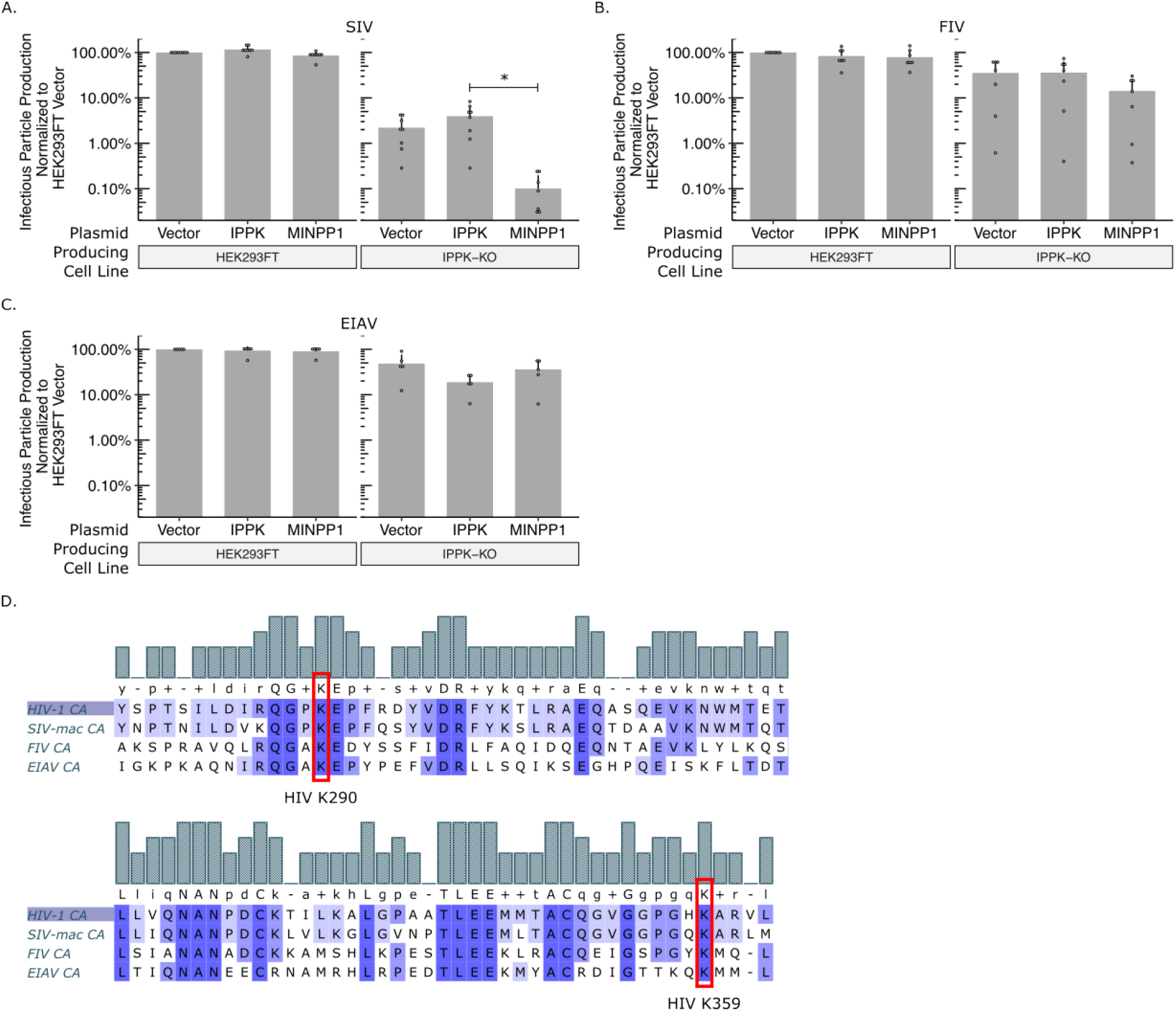
IP6 and IP5 are required assembly co-factors for primate lentiviruses. Bar charts of percent infectious particle release of lentiviruses normalized to virus from HEK293FT cells expressing the empty vector. (A) Simian immunodeficiency virus from macaques (SIV, n = 8). (B) Feline immunodeficiency virus (FIV, n = 7). (C) Equine infectious anemia virus (EIAV, n = 5). (D) Multiple sequence alignment of CA proteins of HIV-1, SIV, FIV, and EIAV. Note the homology of K290 and K359 in HIV-1 to lysines in SIV, FIV, and EIAV.

We next wanted to determine if the sensitivity observed in infectious particle production is reflected in an assembly assay *in vitro* with purified proteins. We have reported previously that IP6 stimulates assembly of EIAV particles, despite the lack of dependence in infectivity assays [40]. We therefore chose to test the stimulation of HIV-1, SIV, FIV, and EIAV in *in vitro* assembly reactions at pH8 and different IP6 concentrations (Fig 9). As expected, addition of as little as 5 μM IP6 stimulated robust assembly of immature, spherical virus like particles (VLPs) for HIV-1 and SIV-mac (Fig 9A-B and E). However, FIV and EIAV required higher concentrations of IP6 and showed more moderate effects (Fig 9C-E), consistent with our previous findings [40]. Interestingly, the construct used for EIAV assembly in the absence of IP6 predominantly forms narrow tubes, which we previously showed to be immature-like lattices [40]. However, in the presence of IP6, they formed predominantly spherical VLPs (Fig 9D and F). Together, these data suggest that IP6 is a requirement for primate lentiviruses and likely acts as an enhancer to promote non-primate lentivirus assembly.

**Fig 9.**
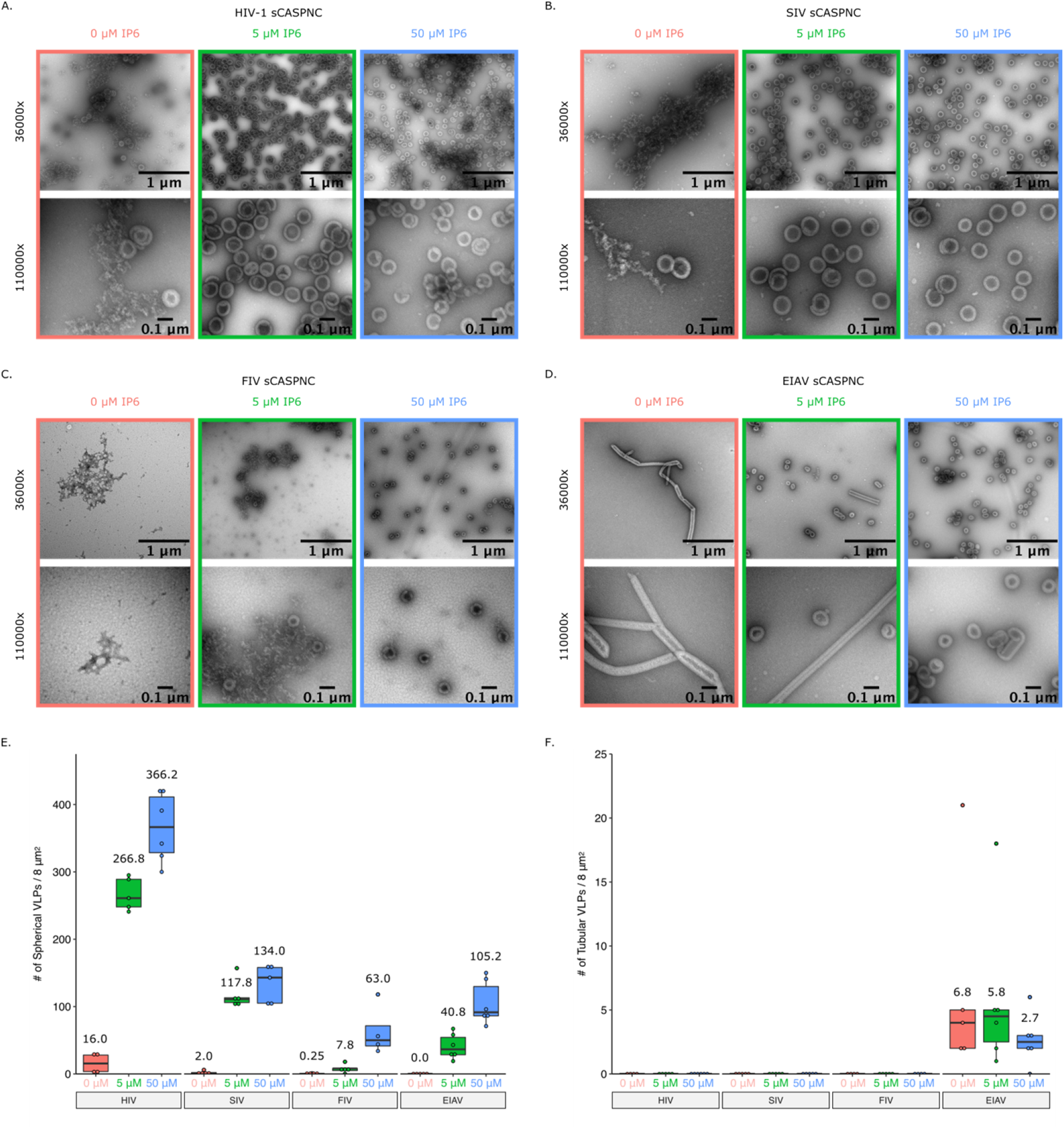
Addition of IP6 stimulates in vitro immature assembly of lentiviruses. Representative images at 36000x and 110000x of virus like particles (VLPs) from *in vitro* assembly reactions at pH8. Assembly reactions were performed with 0, 5, or 50 μM of IP6. (A) VLPs from HIV-1 sCASPNC (ectopic Serine preceding CASPNC) assemblies. (B) VLPs from SIV sCASPNC assemblies. (C) VLPs from FIV sCACSPNC assemblies. (D) VLPs from EIAV sCASPNC assemblies. (E) Quantification of spherical VLPs from each virus assembly reaction (n = 4-6). (F) Quantification of tubular VLPs from each virus assembly reaction (n = 5-6). Should include a definition of the box plot.

## Discussion

### The requirement of IP6 and IP5 for HIV-1 assembly in vivo

IP6 has been shown to be an HIV-1 assembly co-factor. The importance of IP6 has been described in HIV-1 assembly, where it promotes both immature Gag and mature CA assembly, as well as during viral entry, where it stabilizes the capsid en route to the nucleus [10,29,30]. Recently, Mallery *et al.* demonstrated that IP5 is incorporated into viral particles from cells deficient in IP6 production [31]. While release of virus from their IPPK-KO cells was severely diminished, the infectivity of the limited virus that was released was not reduced [31]. This recapitulated the severe loss in virus particle release found in the IPPK-KO cells of Dick *et al.* [10]. Similarly, production of virus in IPMK-KO cells demonstrated that HIV-1 virus particles packaged IP6 despite depletion of cellular IP6 and IP5 levels [31]. While these studies show a role for IP6 and IP5, the absolute requirement of these small molecules had not been addressed.

### Knock-out of IPPK and IPMK affects cellular levels of IP6 and IP5 resulting in loss of virus production

We confirmed the ablation of IP6 pools in the IPPK-KO cell line used in Dick *et al.* [10]. There were, however, slightly elevated levels of IP5 in these cells. Since IP5 can substitute for IP6 *in vitro*, the elevated IP5 could explain the residual virus output from the IPPK-KO cells. Alternatively, IPPK and IP6 have been shown to play roles in many cellular pathways that may have negative effects on virus production. For example, IP6 is involved in the activation of histone deacetylase-1 and mRNA export, with IPPK knock-down causing G1/S phase arrest [41,42]. Our IPPK-KO cells do proliferate more slowly than HEK293FT cells, in concordance with these data. Thus, ablated IP6 and cell arrest could then negatively impact HIV-1 transcriptional regulation, by maintenance of acetylated histones or by block of genome export. Conversely, IP6 can promote necroptosis by directly binding mixed lineage kinase domain-like (MLKL) allowing for plasma membrane rupture [22]. IP6 ablation in IPPK-KO cells prevented IP6 direct binding and activation of MLKL, thereby inhibiting necroptosis. Since HIV-1 infection has been shown to mediate necroptosis [43,44], ablation of IP6 may push infected cells toward apoptosis and induce cytopathic effects in neighboring HIV-1 infected cells, thus, reducing overall infectious particle production in the population of cells.

In our attempt to identify IP5’s role in immature assembly, we knocked-out IPMK using a single guide RNA. The resulting IPMK-KO cell line had residual levels of IP6 and IP5, which correlated with an intermediate loss of virus production. Additionally, residual levels of IP6 and IP5 pointed to an alternative pathway for IP5 synthesis. The role of ITPK1 in inositol phosphate metabolism has not been fully resolved [12,16,21–27]. Since ITPK1 has 5- and 6-kinase activity, this enzyme may compensate for the loss of IPMK (Fig 1A). Attempts to produce an IPMK-ITPK1 double knockout cells were not successful likely because of lethality. This result led us to take the alternative approach of removing residual levels of IP5 and IP6, instead of preventing their biosynthesis.

### Transient removal of IP6 and IP5 ablates HIV-1 infectious particle production

Transient expression of MINPP1 resulted in substantial loss of IP5 in the IPPK-KO cells. This loss of IP5 correlated with a further decrease in release of infectious virus. The inability of IP6- and IP5-depleted cells to release virus is in agreement with the findings of Mallery *et al* [31] and provides evidence for an absolute requirement for these inositol phosphates in immature virus assembly. While MINPP1 is known to remove only the phosphate at the 3 position on the inositol ring, this position is on the equatorial plane of myo-inositol (Fig 3A) [36–38]. Furthermore, removal of 3-phosphates results in dead-end inositol phosphate species according to the currently known metabolism pathway [12]. These inositol phosphate species are currently not known to be re-phosphorylated to produce relevant IP6 and IP5 for HIV-1 assembly [12]. The negative charge on the equatorial plane is critical for coordinating the lysine ring of the MHR K290 in HIV-1 as demonstrated by the low number of VLPs in assembly reactions with IP4 [10]. It is likely that the residual IP5 detected with MINPP1 addition to IPPK-KO cells corresponds to IP5 species that have equatorial hydroxyls and are not efficiently utilized by lentivirus assembly (Fig 3A).

### Depletion of IP6 and IP5 in target cells does not affect susceptibility to infection

Mature HIV-1 virus particles use IP6 to stabilize the Fullerene cone capsid structure. IP6 also has been implicated in hexamer pore interactions with dNTPs, required for reverse transcription, and for trafficking to the nuclear envelope [10,29,30,39]. Therefore, it seemed possible that IP6 and IP5 levels could affect these viral interactions during viral entry, and thus cell susceptibility to infection. Here, we demonstrated cells depleted of IP6 and IP5 are just as susceptible to infection as HEK293FT cells. Our data suggest that if IP6 or IP5 are required for viral entry and trafficking, the molecules incorporated during viral assembly are sufficient for this process. We speculate that while cells devoid of IP6 and IP5 cannot produce infectious HIV-1 virus particles, mutants in Gag might be able to do so. Such mutants might be used to address the dynamics of IP6-capsid interactions at early stages of infection. Furthermore, since inositol phosphates have been implicated in immune responses such as RIG-I signaling [45], there may be signaling responses that can affect the rate of infection. More detailed kinetic studies would be required to investigate this possibility.

### Requirement of IP6 and IP5 is conserved across primate lentiviruses and likely acts as an enhancer for assembly of non-primate lentiviruses

The robust requirement of IP6 and IP5 is conserved across primate lentiviral species. The lower Gag amino acid sequence homology between lentiviruses and retroviruses of other genera correlates with the lack of a phenotype for beta- and gamma-retroviruses in cells with ablated IP6 and IP5 levels. This suggests that either their structural proteins have relatively stable hexagonal lattice structures and do not require a coordinating molecule, or that another small molecule coordinates structural protein assembly. How viruses evolved to use IP6 is still a topic of great interest and should be further studied.

## Conclusion

In this study, we present data to show that IP6 and IP5 are required for HIV-1 infectious virus particle assembly. Additionally, this robust requirement is likely conserved across primate lentiviruses, but not for other retrovirus genera. While IP6 at high molar concentrations can stimulate *in vitro* assembly for non-primate lentiviruses, the physiological relevance remains to be determined. Additionally, understanding at what point IP6 is incorporated into the forming Gag lattice in the cell, for example nucleating assembly of Gag hexamers, may provide new targets for therapeutics.

## Materials and Methods

### Plasmid constructs

All lentiviral vectors for CRISPR/Cas-9 delivery were pseudotyped with VSV-g (NIH AIDS Reagent Program) [46]. CRISPR/Cas-9 vectors were derived from the plasmid lentiCRISPRv2 (a gift from Feng Zhang; Addgene plasmid # 52961; http://n2t.net/addgene:52961; RRID:Addgene_52961) [32]. Guide sequences for *IPPK* and *IPMK* were obtained from the Human GeCKOv2 CRISPR knockout pooled libraries (a gift from Feng Zhang; Addgene #1000000048, #1000000049) [32]. Briefly, nucleotide bases as per the lentCRISPRv2 protocol were added to the specific sequences for *IPPK*, *IPMK*, and *ITPK* from the pool, and were cloned into the lentiCRISPRv2 plasmid (Table 1) [32]. The CRISPR/Cas9 vector with guide sequences were delivered via the packaging vector psPAX2 (a gift from Didier Trono; Addgene plasmid # 12260; http://n2t.net/addgene:12260; RRID:Addgene_12260).

**Table 1.**
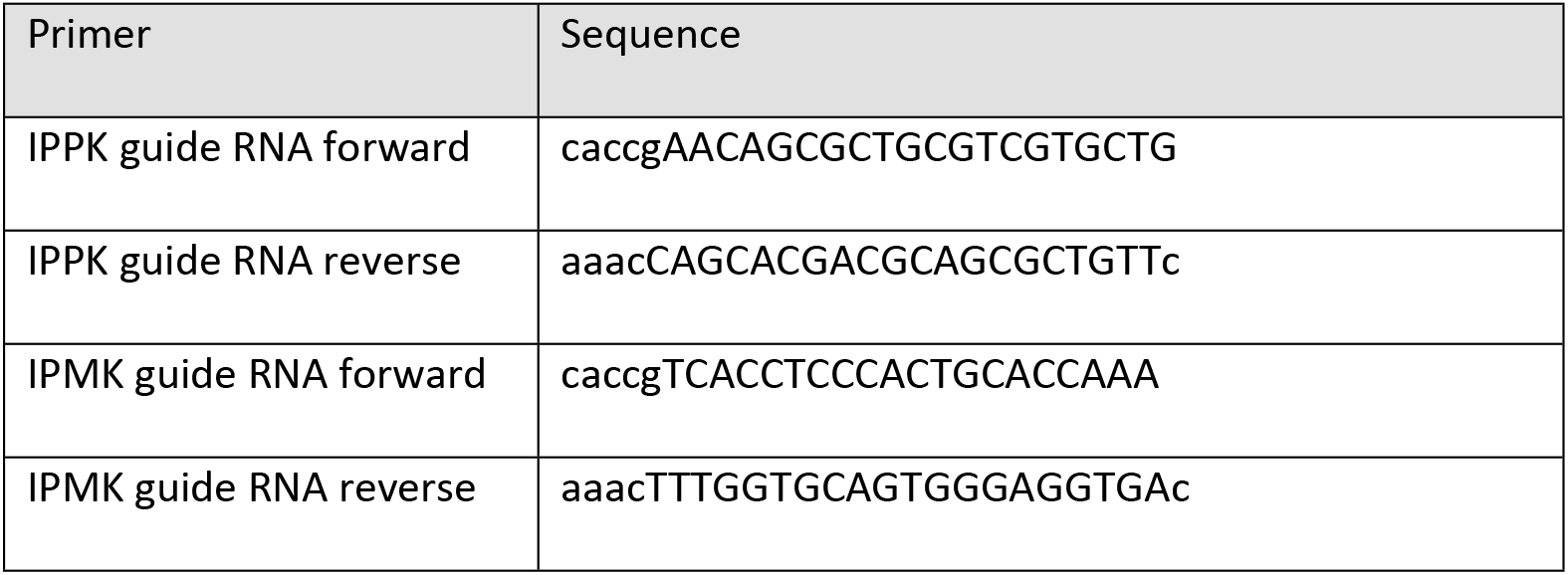
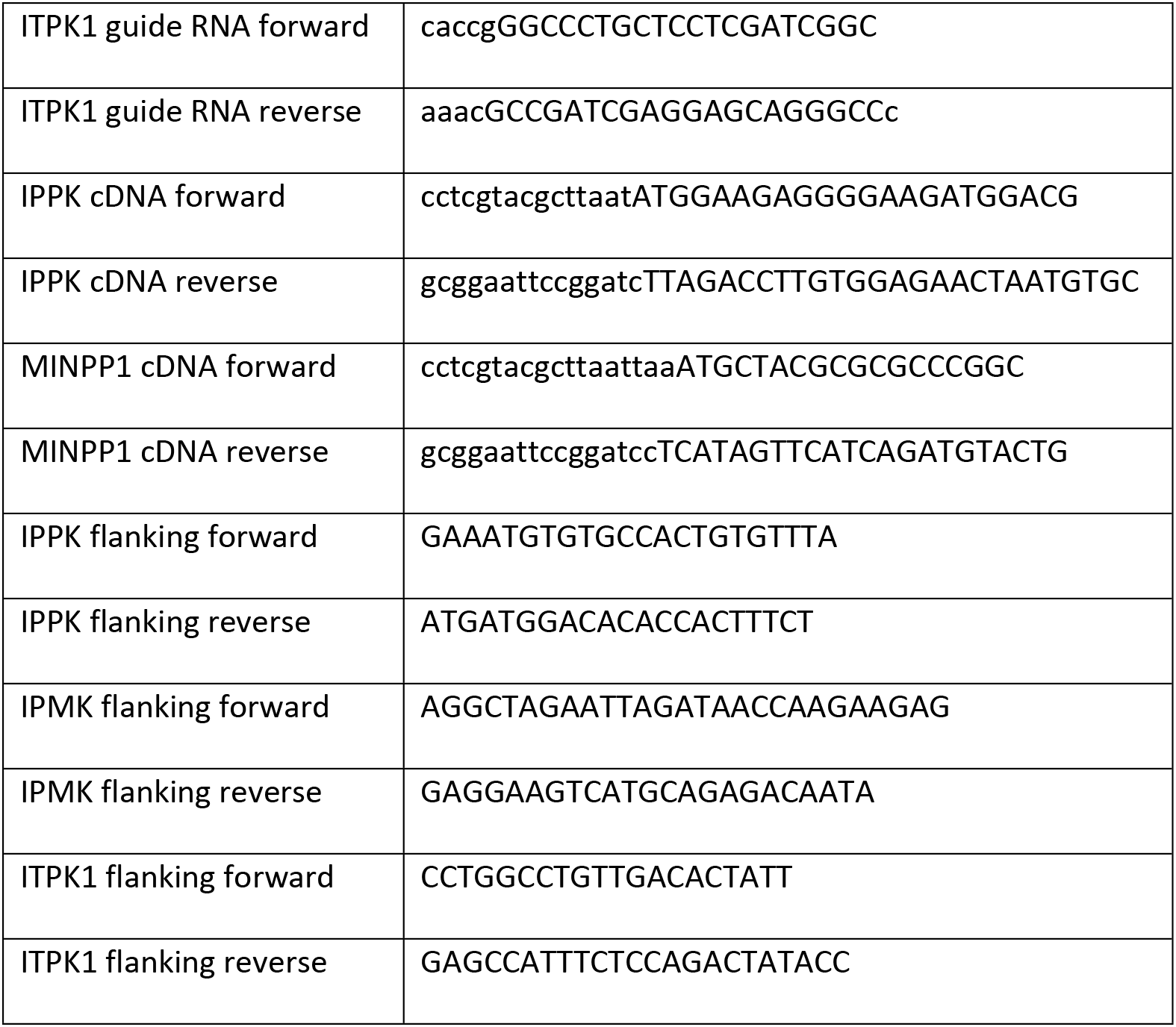
Primers used for cloning.

All cDNA vectors were packaged with a CMV-MLV-Gag-Pol expression plasmid (kindly provided by Walther Mothes, Yale University) pseudotyped with VSVg (NIH AIDS Reagent Program) [46]. Primers for cDNA amplification were ordered from IDTDNA (Table 1). Primers consisted of 3’ sequences matching the coding sequences for *IPPK* and *MINPP1* and had 15 base pair overhangs for InFusion cloning (Clonetech, PT3669-5; Cat. No. 631516) into expression plasmid pQCXIH (Clonetech) using the restriction sites PacI and BamHI. The MINPP1 IRES GFP was made by replacing the hygromycin resistance cassette from the previous clone with IRES-GFP via InFusion cloning.

All retroviruses were pseudotyped with VSV-g (NIH AIDS Reagent Program) [46]. HIV-1 ^ΔEnv^ consisted of NL4-3-derived proviral vector with a 3’ cytomegalovirus (CMV) driven green fluorescent protein (GFP) and defective for Vif, Vpr, Nef, & Env (kindly provided by Vineet Kewal-Ramani, National Cancer Institute-Frederick). HIV-1-CD4 was made by replacing GFP from the previous clone with CD4 via InFusion cloning. SIV^ΔEnv^ consisted of the Gag-Pol expression plasmid pUpSVOΔΫ and the reporter vector plasmid pV1eGFPSVO (kindly provided by Hung Fan, University of California-Irvine). FIV ^ΔEnv^ consisted of the Gag-Pol expression plasmid pFP93 and the reporter vector plasmid pGinsin (kindly provided by Eric Poeschla, University of Colorado-Denver) [47]. EIAV ^ΔEnv^ consisted of the Gag-Pol expression plasmid pONY3.1 and the reporter vector plasmid pONY8.0-GFP (kindly provided by Nicholas D. Mazarakis, Imperial College-London) [48]. MPMV ^ΔEnv^ consisted of an Env deficient expression plasmid pSARM into which our lab engineered a CMV driven GFP reporter before the 3’ LTR (kindly provided by Eric Hunter, Emory University). MLV ^ΔEnv^ consisted of the CMV-MLV-Gag-Pol expression plasmid (kindly provided by Walther Mothes, Yale University) and the reporter vector plasmid pQCXIP-GFP (a gift from Michael Grusch (Addgene plasmid # 73014; http://n2t.net/addgene:73014; RRID:Addgene_73014)).

### Cells and knock-out of cellular genes

The HEK293FT cell line was obtained from Invitrogen and maintained in Dulbecco’s modified Eagle’s medium (DMEM) supplemented with 10% fetal bovine serum (FBS), 2 mM glutamine, 1 mM sodium pyruvate, 10 mM nonessential amino acids, and 1% minimal essential medium vitamins. Knock-out cell lines were obtained via transduction of HEK293FT cells with lentiCRISPRv2 containing the guide sequences for *IPPK* and *IPMK* genes (Table 1). 48 hrs post transduction culture media was replaced with media containing 1 μg/mL puromycin and incubated till complete death of non-transduced control cells (~48-72 hrs). Clonal isolates were obtained by sparse plating of surviving cells on a 10 cm dish and allowing colonies to grow. Colonies were then picked and expanded in new plates. KO was verified by amplification of genomic DNA flanking the CRISPR target site (Table 1), direct sequencing of the PCR product, and sequence analysis.

### Virus production and transductions

HIV (NL4-3 derived), SIV (mac), EIAV (pony), MLV (maloney), VLPs were produced by polyethylenimine (PEI, made in house) transfection of HEK293FTs or KO derivatives at 50% confluence with 900 ng of viral plasmids plus VSV-g in a 10:1 ratio [49]. Media containing virus (viral media) was collected two days post transfection. Viral media was then frozen at −80°C for a minimum of 1 hr to lyse cells, thawed in a 37°C water bath, precleared by centrifugation at 3000 x g for 5 min and supernatant collected. Aliquots after titration were stored at −80°C and subsequently used for assays.

Viral media was titered on HEK293FT cells by serial dilution. Viral media was then added to fresh HEK293FT cells at low MOI to prevent infection saturation. Infected cells were collected and assayed via flow cytometry. Infections were then normalized to percent of infections in WT HEK293FT cells and presented as relative particle production.

### Separation of inositol phosphates

Inositol phosphates were separated and quantified as per Wilson *et al.* [35]. All steps were performed on ice and 100 μL of 100 μM IP6 and 100 μL of 10 μM IP6 standards were treated in parallel as a control. Briefly, inositol phosphates were extracted from counted cells (HEK293FT and IPPK-KO) by suspension in 1 M perchloric acid. Cells were pelleted and supernatants were transferred to tubes containing 4 mg of TiO_2_ beads (GL Sciences Inc., Titansphere; Cat. No. 5020-75000) in 50 μL 1 M perchloric acid to bind inositol phosphates. Washed inositol phosphates bound to TiO_2_ beads were eluted with 10% ammonium hydroxide, beads pelleted, and supernatant collected. Supernatant was then concentrated to 10uL and pH neutralized by SpeedVac centrifugation.

A 33% PAGE large gel (16 x 20 cm; TBE pH 8, Acrylamide:Bis 19:1, SDS) was cast and pre-run for 30 min at 500 V. The entirety of concentrated samples (10 μL) was mixed with 50 μL bromophenol blue loading buffer (6X Buffer). The samples were then run over night at 4°C at 1000 V until the loading dye had traveled through one-third of the gel. The gel was then stained with toluidine blue for 30 min and destained. Gels were then imaged. Molar concentrations were calculated assuming a cell diameter of 15 μM [50]

### Surface labeling of cells

Cells were washed with PBS and treated with 10 mM EDTA. Cells were then collected with PBS, centrifuged at 300 x g for 5 min, and supernatant removed. Cells were blocked with 5% goat serum in PBS for 30 min on ice. Cells were centrifuged at 300 x g for 5 min and supernatant removed. Anti-CD4 antibody conjugated to Alexa-Fluor 555 was applied to cells at 1:100 in 1% goat serum in PBS for 1 hr. Cells were then washed 3 x with PBS, suspended in 400 μL of PBS, and 100 μL of 10% paraformaldehyde (PFA) added to fix cells. After incubation for 20 min on ice, cells were centrifuged at 300 x g for 5 min and supernatant was removed. Cells were resuspended in 200 μL of PBS and analyzed via flow cytometry.

### Flow cytometry

Cells in 6- and 12-well format were washed with PBS and treated with 10 mM TrypLE™ Express Enzyme (Gibco; Cat. No. 12605028). Cells were then collected with PBS and added to 10% PFA to a final concentration of 4%. After 10-20 min incubation at room temperature, the cells were centrifuged at 300 x g for 5 min, supernatant removed, and 300 μL of PBS added. Cells were analyzed for fluorescence using an Accuri C6 flow cytometer.

### Western blot

Supernatant collected after preclearing thawed media containing virus was pelleted via centrifugation through a 20% sucrose cushion (20% sucrose, 100 mM NaCl, 10 mM Tris, 1 mM EDTA, pH 7.5) for 2 h at 30000 x g at 4°C. Supernatant and sucrose buffer were aspirated off, leaving a small amount of sucrose buffer so as not to aspirate the viral pellet (~10 μL). Ten μL of 2x sample buffer (50 mM Tris, 2% sodium dodecyl sulfate [SDS], 20% glycerol, 5% β-mercaptoethanol) was added to pelleted virus and heated to 95°C for 5 min before loading.

Cell samples were washed with PBS and trypsinized with 10 mM EDTA. Cells were then collected with PBS, centrifuged at 300 x g for 5 min, and supernatant removed. Twenty μL of RIPA extraction buffer with protease inhibitor was then added to each sample [51]. The samples were then kept on ice and vortexed every 5 min for 20 min, followed by centrifugation at 10000 rpm for 10 min at 4°C. Supernatant was then transferred to a new tube, 20 μL of 2x sample buffer added, and heated to 95°C for 5 min before loading.

Samples were separated on a 10% SDS-PAGE gel, and transferred onto a 0.22 μm pore size polyvinylidene difluoride (PVDF) membrane. Membranes were blocked for 1 hr at room temperature with 5% nonfat dry milk in PBS-tween. Membranes were then incubated with anti-HIV p24 hybridoma medium was diluted 1:500 (HIV-1 p24 hybridoma [183-H12-5C], obtained from NIH AIDS Reagent Program) from Bruce Chesebro [52] for 1 hr at room temperature. After blots were washed with PBST (3 x for 5 min), horseradish peroxidase (HRP)-conjugated secondary antibody was applied at 1:10,000 to all blots. After 1 hr, blots were again washed 3x with PBST and imaged. Horseradish peroxidase-linked anti-mouse (A5278), was obtained from Sigma. Luminata Classico Western HRP substrate (Millipore) was used for visualization of the membranes with a chemiluminescence image analyzer (UVP BioSpectrum 815 Imaging System).

### Protein purification and in vitro assembly of CANC protein

Protein purification, *in vitro* assembly, and imaging of all lentiviral CANC proteins was performed as previously described in Dick *et al.* [40] and briefly here. 50 μM protein purified from bacteria was mixed with 10 μM GT25 oligo without or with 5 μM or 10 μM IP6. 30 μL assembly reactions were dialyzed against 2 mL buffer (50 mM Tris-HCl pH 8, 100 mM NaCl, 2 mM TCEP, without or with 5 μM or 10 μM IP6. 4 hrs post dialysis, assembly reactions were adjusted to 200 μL, spotted onto formvar/carbon grids, stained with 2% uranly acetate, and imaged on a FEI Morgagni transmission electron microscope.

### Data analysis

Flow cytometry data was analyzed using FlowJo™ software [53]. Values for fluorescence were exported to Excel spreadsheet. Images were analyzed using Fiji (ImageJ) [54]. Western blot images were converted to 8-bit, Fiji’s gel analysis tools used to calculate density, and values exported to an Excel spreadsheet. VLPs from electron micrographs were quantified manually using Fiji’s cell counting tool and values recorded in an Excel spreadsheet. Values from Excel spreadsheets were formatted for statistical analysis via R [55] and exported to CSV format. RStudio was used to analyze data and create figures [56]. UGene was used for plasmid cloning, sequence analysis, and chromatogram image generation [57]. Final figures were prepared using Inkscape [58].

## Acknowledgments

This work was supported by the National Institute of Allergy and Infectious Diseases (NIAID) under awards R21AI143363 to Marc C. Johnson, R01AI147890 to Robert A. Dick, and R01AI150454 to Volker M. Vogt. We thank John York for helpful discussions.

## Author Contributions

Conceptualization: Marc C Johnson, Clifton L. Ricana

Data Curation: Clifton L Ricana, Marc C Johnson

Formal Analysis: Clifton L Ricana, Marc C Johnson

Funding Acquisition: Marc C Johnson

Investigation: Clifton L Ricana, Terri D Lyddon, Robert A Dick

Methodology: Clifton L Ricana, Marc C Johnson

Project Administration: Marc C Johnson, Clifton L Ricana

Resources: Marc C Johnson, Robert A Dick

Supervision: Marc C Johnson

Validation: Clifton L Ricana, Terri D Lyddon, Robert A Dick, Marc C Johnson

Visualization: Clifton L Ricana, Robert A Dick

Writing – Original Draft Preparation: Clifton L Ricana, Marc C Johnson

Writing – Review & Editing: Clifton L Ricana, Marc C Johnson, Robert A Dick

## References

1. Freed EO. HIV-1 assembly, release and maturation. Nat Rev Microbiol. 2015;13: 484–496. doi:10.1038/nrmicro3490

2. Gross I, Hohenberg H, Wilk T, Wiegers K, Grättinger M, Müller B, et al. A conformational switch controlling HIV-1 morphogenesis. EMBO J. 2000;19: 103–113. doi:10.1093/emboj/19.1.103

3. Datta SAK, Zhao Z, Clark PK, Tarasov S, Alexandratos JN, Campbell SJ, et al. Interactions between HIV-1 Gag molecules in solution: an inositol phosphate-mediated switch. J Mol Biol. 2007;365: 799–811. doi:10.1016/j.jmb.2006.10.072

4. Schur FKM, Obr M, Hagen WJH, Wan W, Jakobi AJ, Kirkpatrick JM, et al. An atomic model of HIV-1 capsid-SP1 reveals structures regulating assembly and maturation. Science. 2016;353: 506–508. doi:10.1126/science.aaf9620

5. Wagner JM, Zadrozny KK, Chrustowicz J, Purdy MD, Yeager M, Ganser-Pornillos BK, et al. Crystal structure of an HIV assembly and maturation switch. Elife. 2016;5. doi:10.7554/eLife.17063

6. Wlodawer A, Erickson JW. Structure-based inhibitors of HIV-1 protease. Annu Rev Biochem. 1993;62: 543–585. doi:10.1146/annurev.bi.62.070193.002551

7. Keller PW, Huang RK, England MR, Waki K, Cheng N, Heymann JB, et al. A two-pronged structural analysis of retroviral maturation indicates that core formation proceeds by a disassembly-reassembly pathway rather than a displacive transition. J Virol. 2013;87: 13655–13664. doi:10.1128/JVI.01408-13

8. Campbell S, Fisher RJ, Towler EM, Fox S, Issaq HJ, Wolfe T, et al. Modulation of HIV-like particle assembly in vitro by inositol phosphates. Proc Natl Acad Sci USA. 2001;98: 10875–10879. doi:10.1073/pnas.191224698

9. Munro JB, Nath A, Färber M, Datta SAK, Rein A, Rhoades E, et al. A conformational transition observed in single HIV-1 Gag molecules during in vitro assembly of virus-like particles. J Virol. 2014;88: 3577–3585. doi:10.1128/JVI.03353-13

10. Dick RA, Zadrozny KK, Xu C, Schur FKM, Lyddon TD, Ricana CL, et al. Inositol phosphates are assembly co-factors for HIV-1. Nature. 2018;560: 509–512. doi:10.1038/s41586-018-0396-4

11. Letcher AJ, Schell MJ, Irvine RF. Do mammals make all their own inositol hexakisphosphate? Biochem J. 2008;416: 263–270. doi:10.1042/BJ20081417

12. Reactome Team, Anwesha Bohler, Martina Kutmon. Inositol phosphate metabolism (Homo sapiens) - WikiPathways. 1 Nov 2018 [cited 10 Mar 2020]. Available: https://www.wikipathways.org/index.php/Pathway:WP2741

13. Ives EB, Nichols J, Wente SR, York JD. Biochemical and functional characterization of inositol 1,3,4,5, 6-pentakisphosphate 2-kinases. J Biol Chem. 2000;275: 36575–36583. doi:10.1074/jbc.M007586200

14. Verbsky JW, Wilson MP, Kisseleva MV, Majerus PW, Wente SR. The synthesis of inositol hexakisphosphate. Characterization of human inositol 1,3,4,5,6-pentakisphosphate 2-kinase. J Biol Chem. 2002;277: 31857–31862. doi:10.1074/jbc.M205682200

15. Verbsky J, Lavine K, Majerus PW. Disruption of the mouse inositol 1,3,4,5,6-pentakisphosphate 2-kinase gene, associated lethality, and tissue distribution of 2-kinase expression. Proc Natl Acad Sci USA. 2005;102: 8448–8453. doi:10.1073/pnas.0503656102

16. Monserrate JP, York JD. Inositol phosphate synthesis and the nuclear processes they affect. Curr Opin Cell Biol. 2010;22: 365–373. doi:10.1016/j.ceb.2010.03.006

17. Frederick JP, Mattiske D, Wofford JA, Megosh LC, Drake LY, Chiou S-T, et al. An essential role for an inositol polyphosphate multikinase, Ipk2, in mouse embryogenesis and second messenger production. Proc Natl Acad Sci USA. 2005;102: 8454–8459. doi:10.1073/pnas.0503706102

18. Yang X, Shears SB. Multitasking in signal transduction by a promiscuous human Ins(3,4,5,6)P(4) 1-kinase/Ins(1,3,4)P(3) 5/6-kinase. Biochem J. 2000;351 Pt 3: 551–555.

19. Malabanan MM, Blind RD. Inositol polyphosphate multikinase (IPMK) in transcriptional regulation and nuclear inositide metabolism. Biochem Soc Trans. 2016;44: 279–285. doi:10.1042/BST20150225

20. Kim E, Ahn H, Kim MG, Lee H, Kim S. The Expanding Significance of Inositol Polyphosphate Multikinase as a Signaling Hub. Mol Cells. 2017;40: 315–321. doi:10.14348/molcells.2017.0066

21. Dovey CM, Diep J, Clarke BP, Hale AT, McNamara DE, Guo H, et al. MLKL Requires the Inositol Phosphate Code to Execute Necroptosis. Mol Cell. 2018;70: 936–948.e7. doi:10.1016/j.molcel.2018.05.010

22. McNamara DE, Dovey CM, Hale AT, Quarato G, Grace CR, Guibao CD, et al. Direct Activation of Human MLKL by a Select Repertoire of Inositol Phosphate Metabolites. Cell Chem Biol. 2019;26: 863–877.e7. doi:10.1016/j.chembiol.2019.03.010

23. Riley AM, Deleu S, Qian X, Mitchell J, Chung S-K, Adelt S, et al. On the contribution of stereochemistry to human ITPK1 specificity: Ins(1,4,5,6)P4 is not a physiologic substrate. FEBS Lett. 2006;580: 324–330. doi:10.1016/j.febslet.2005.12.016

24. Chamberlain PP, Qian X, Stiles AR, Cho J, Jones DH, Lesley SA, et al. Integration of inositol phosphate signaling pathways via human ITPK1. J Biol Chem. 2007;282: 28117–28125. doi:10.1074/jbc.M703121200

25. Saiardi A, Cockcroft S. Human ITPK1: a reversible inositol phosphate kinase/phosphatase that links receptor-dependent phospholipase C to Ca2+-activated chloride channels. Sci Signal. 2008;1: pe5. doi:10.1126/stke.14pe5

26. Shears SB. Molecular basis for the integration of inositol phosphate signaling pathways via human ITPK1. Adv Enzyme Regul. 2009;49: 87–96. doi:10.1016/j.advenzreg.2008.12.008

27. Desfougères Y, Wilson MSC, Laha D, Miller GJ, Saiardi A. ITPK1 mediates the lipid-independent synthesis of inositol phosphates controlled by metabolism. PNAS. 2019 [cited 21 Nov 2019]. doi:10.1073/pnas.1911431116

28. Wundenberg T, Grabinski N, Lin H, Mayr GW. Discovery of InsP6-kinases as InsP6-dephosphorylating enzymes provides a new mechanism of cytosolic InsP6 degradation driven by the cellular ATP/ADP ratio. Biochem J. 2014;462: 173–184. doi:10.1042/BJ20130992

29. Mallery DL, Márquez CL, McEwan WA, Dickson CF, Jacques DA, Anandapadamanaban M, et al. IP6 is an HIV pocket factor that prevents capsid collapse and promotes DNA synthesis. Elife. 2018;7. doi:10.7554/eLife.35335

30. Dick RA, Mallery DL, Vogt VM, James LC. IP6 Regulation of HIV Capsid Assembly, Stability, and Uncoating. Viruses. 2018;10. doi:10.3390/v10110640

31. Mallery DL, Faysal KMR, Kleinpeter A, Wilson MSC, Vaysburd M, Fletcher AJ, et al. Cellular IP6 Levels Limit HIV Production while Viruses that Cannot Efficiently Package IP6 Are Attenuated for Infection and Replication. Cell Reports. 2019;29: 3983–3996.e4. doi:10.1016/j.celrep.2019.11.050

32. Sanjana NE, Shalem O, Zhang F. Improved vectors and genome-wide libraries for CRISPR screening. Nat Methods. 2014;11: 783–784. doi:10.1038/nmeth.3047

33. Shalem O, Sanjana NE, Hartenian E, Shi X, Scott DA, Mikkelson T, et al. Genome-scale CRISPR-Cas9 knockout screening in human cells. Science. 2014;343: 84–87. doi:10.1126/science.1247005

34. Losito O, Szijgyarto Z, Resnick AC, Saiardi A. Inositol pyrophosphates and their unique metabolic complexity: analysis by gel electrophoresis. PLoS ONE. 2009;4: e5580. doi:10.1371/journal.pone.0005580

35. Wilson MSC, Bulley SJ, Pisani F, Irvine RF, Saiardi A. A novel method for the purification of inositol phosphates from biological samples reveals that no phytate is present in human plasma or urine. Open Biol. 2015;5: 150014. doi:10.1098/rsob.150014

36. Chi H, Yang X, Kingsley PD, O’Keefe RJ, Puzas JE, Rosier RN, et al. Targeted deletion of Minpp1 provides new insight into the activity of multiple inositol polyphosphate phosphatase in vivo. Mol Cell Biol. 2000;20: 6496–6507. doi:10.1128/mcb.20.17.6496-6507.2000

37. Windhorst S, Lin H, Blechner C, Fanick W, Brandt L, Brehm MA, et al. Tumour cells can employ extracellular Ins(1,2,3,4,5,6)P(6) and multiple inositol-polyphosphate phosphatase 1 (MINPP1) dephosphorylation to improve their proliferation. Biochem J. 2013;450: 115–125. doi:10.1042/BJ20121524

38. Kilaparty SP, Agarwal R, Singh P, Kannan K, Ali N. Endoplasmic reticulum stress-induced apoptosis accompanies enhanced expression of multiple inositol polyphosphate phosphatase 1 (Minpp1): a possible role for Minpp1 in cellular stress response. Cell Stress Chaperones. 2016;21: 593–608. doi:10.1007/s12192-016-0684-6

39. Huang P-T, Summers BJ, Xu C, Perilla JR, Malikov V, Naghavi MH, et al. FEZ1 Is Recruited to a Conserved Cofactor Site on Capsid to Promote HIV-1 Trafficking. Cell Rep. 2019;28: 2373–2385.e7. doi:10.1016/j.celrep.2019.07.079

40. Dick RA, Xu C, Morado DR, Kravchuk V, Ricana CL, Lyddon TD, et al. Structures of immature EIAV Gag lattices reveal a conserved role for IP6 in lentivirus assembly. PLOS Pathogens. 2020;16: e1008277. doi:10.1371/journal.ppat.1008277

41. Okamura M, Yamanaka Y, Shigemoto M, Kitadani Y, Kobayashi Y, Kambe T, et al. Depletion of mRNA export regulator DBP5/DDX19, GLE1 or IPPK that is a key enzyme for the production of IP6, resulting in differentially altered cytoplasmic mRNA expression and specific cell defect. PLoS ONE. 2018;13: e0197165. doi:10.1371/journal.pone.0197165

42. Watson PJ, Millard CJ, Riley AM, Robertson NS, Wright LC, Godage HY, et al. Insights into the activation mechanism of class I HDAC complexes by inositol phosphates. Nat Commun. 2016;7: 11262. doi:10.1038/ncomms11262

43. Pan T, Wu S, He X, Luo H, Zhang Y, Fan M, et al. Necroptosis takes place in human immunodeficiency virus type-1 (HIV-1)-infected CD4+ T lymphocytes. PLoS ONE. 2014;9: e93944. doi:10.1371/journal.pone.0093944

44. Gaiha GD, McKim KJ, Woods M, Pertel T, Rohrbach J, Barteneva N, et al. Dysfunctional HIV-specific CD8+ T cell proliferation is associated with increased caspase-8 activity and mediated by necroptosis. Immunity. 2014;41: 1001–1012. doi:10.1016/j.immuni.2014.12.011

45. Pulloor NK, Nair S, McCaffrey K, Kostic AD, Bist P, Weaver JD, et al. Human genome-wide RNAi screen identifies an essential role for inositol pyrophosphates in Type-I interferon response. PLoS Pathog. 2014;10: e1003981. doi:10.1371/journal.ppat.1003981

46. Chang L-J, Urlacher V, Iwakuma T, Cui Y, Zucali J. Efficacy and safety analyses of a recombinant human immunodeficiency virus type 1 derived vector system. Gene Ther. 1999;6: 715–728. doi:10.1038/sj.gt.3300895

47. Saenz DT, Barraza R, Loewen N, Teo W, Poeschla EM. Feline Immunodeficiency Virus-Based Lentiviral Vectors. Cold Spring Harbor Protocols. 2012;2012: pdb.ip067579-pdb.ip067579. doi:10.1101/pdb.ip067579

48. Mitrophanous K, Yoon S, Rohll J, Patil D, Wilkes F, Kim V, et al. Stable gene transfer to the nervous system using a non-primate lentiviral vector. Gene Ther. 1999;6: 1808–1818. doi:10.1038/sj.gt.3301023

49. Boussif O, Lezoualc’h F, Zanta MA, Mergny MD, Scherman D, Demeneix B, et al. A versatile vector for gene and oligonucleotide transfer into cells in culture and in vivo: polyethylenimine. Proceedings of the National Academy of Sciences. 1995;92: 7297–7301. doi:10.1073/pnas.92.16.7297

50. Milo R, Jorgensen P, Moran U, Weber G, Springer M. BioNumbers—the database of key numbers in molecular and cell biology. Nucleic Acids Research. 2010;38: D750–D753. doi:10.1093/nar/gkp889

51. RIPA buffer. Cold Spring Harb Protoc. 2006;2006: pdb.rec10035. doi:10.1101/pdb.rec10035

52. Chesebro B, Wehrly K, Nishio J, Perryman S. Macrophage-tropic human immunodeficiency virus isolates from different patients exhibit unusual V3 envelope sequence homogeneity in comparison with T-cell-tropic isolates: definition of critical amino acids involved in cell tropism. J Virol. 1992;66: 6547–6554.

53. FlowJo™ software for Mac. Ashland, OR: Becton, Dickinson and Company; 2019. Available: https://www.flowjo.com

54. Schindelin J, Arganda-Carreras I, Frise E, Kaynig V, Longair M, Pietzsch T, et al. Fiji: an open-source platform for biological-image analysis. Nat Methods. 2012;9: 676–682. doi:10.1038/nmeth.2019

55. R Core Team. R: A Language and Environment for Statistical Computing. Vienna, Austria: R Foundation for Statistical Computing; YEAR. Available: https://www.R-project.org

56. RStudio Team. RStudio: Integrated Development Environment for R. Boston, MA: RStudio, Inc.; 2015. Available: http://www.rstudio.com/

57. Okonechnikov K, Golosova O, Fursov M, UGENE team. Unipro UGENE: a unified bioinformatics toolkit. Bioinformatics. 2012;28: 1166–1167. doi:10.1093/bioinformatics/bts091

58. Inkscape. 2020. Available: https://inkscape.org

